# Mechanically Induced Integrin Ligation Mediates Intracellular Calcium Signaling with Single Pulsating Cavitation Bubbles

**DOI:** 10.1101/2020.10.25.353904

**Authors:** Fenfang Li, Tae Hyun Park, George Sankin, Christopher Gilchrist, Defei Liao, Chon U Chan, Zheng Mao, Brenton D. Hoffman, Pei Zhong

## Abstract

Ultrasound or shockwave-induced cavitation is used therapeutically to stimulate neural and muscle tissue, but the mechanisms underlying this mechanotransduction are unclear. Intracellular Ca^2+^ signaling is one of the earliest events in mechanotransduction. In this study, we investigate the mechanism of Ca^2+^ signaling in individual HEK293T cells stimulated by single cavitation bubbles. Ca^2+^ responses are rare at cell-bubble distance that avoids membrane poration, even with overexpression of the mechanosensitive ion channel Piezo1, but could be increased in frequency to 42% of cells by attaching RGD beads to the apical surface of the cells. By using Piezo1 knockout and Piezo1-expressing cells, integrin-blocking antibodies, and inhibitors of P2X ion channels, key molecular players are identified in the RGD bead-enhanced Ca^2+^ response: increased integrin ligation by substrate ECM triggers ATP release and activation of P2X—but not Piezo1—ion channels. These molecular players have not been examined previously in cavitation-induced Ca^2+^ signaling. The resultant Ca^2+^ influx causes dynamic changes in cell spread area. This approach to eliciting a Ca^2+^ response with cavitation microbubbles without cell injury, and the uncovered mechanotransduction mechanism by which increased integrin-ligation mediates ATP release and Ca^2+^ signaling will inform new strategies to stimulate tissues with ultrasound and shockwaves.

## 1. Introduction

Ultrasound can be focused deep in brain or other tissue to allow non-invasive imaging and therapy.^[1, 2]^ Therapeutic applications of ultrasound include shockwave lithotripsy,^[3]^ blood-brain barrier opening,^[4]^ targeted drug and gene delivery,^[5, 6]^ tissue ablation, and induction of anti-tumor immune responses by high-intensity focused ultrasound (HIFU) and histotripsy.^[7–10]^ The biological effects of therapeutic ultrasound are often caused by cavitation bubbles, which induce shear and jetting flow near interfaces,^[11–16^ leading to cell mechanotransduction and intracellular Ca^2+^ signaling. Ultrasound-induced microbubble oscillation is attractive for its potential utility in neuromodulation and cancer immunotherapy.^[17, 18]^ To enhance the intended therapeutic effects of ultrasound treatment, a better understanding of the underlying mechanism is needed.

Cells sense their physical environment through mechanotransduction, the process of converting mechanical forces into biochemical signals that control cell functions. Intracellular Ca^2+^ signaling plays a crucial role in mechanotransduction and regulates myriad cell processes including exocytosis, contraction, transcription, and proliferation. ^[19, 20]^ Ca^2+^ signaling is one of the earliest events in mechanotransduction under quasi-static cell loading,^[21–24]^ and has been widely studied.^[25–27]^ Cavitation microbubbles exert impulsive shear flow and high strain-rate loading on cells and can elicit a Ca^2+^ response, ^[13, 28–30]^ but the mechanism by which this occurs is not well-understood.

Three mechanisms are commonly observed in shear-induced mechanotransduction: 1) direct activation of mechanosensitive ion channels such as Piezo1;^[31]^ 2) triggered release of ATP, which activates purinoreceptors (P2X);^[32]^ and 3) ligation of integrins by extracellular matrix (ECM) proteins.^[33–36]^ The first two mechanisms involve Ca^2+^ influx through ion channels; the third mechanism involves formation of focal adhesions (FAs) on the basal cell surface and integrin-mediated signaling. Quasi-static loading of integrin-bound beads on the apical surfaces of cells can remotely alter basal focal adhesions via integrin-cytoskeleton interactions.^[37]^

High-frequency ultrasound (30–150 MHz) has been shown to directly activate mechanosensitive ion channels without microbubbles.^[17, 38–40]^ However, the tissue penetration depth at such high frequencies is shallow (<5 mm), limiting *in vivo* applications. In contrast, at regular ultrasound frequencies of 1–2 MHz, microbubbles are often required to elicit a Ca^2+^ response, as shown in *C. elegans* and in human mesenchymal stem cells and HEK293T cells.^[17, 18, 41]^

In this work we aim to elucidate the determinants of intracellular Ca^2+^ responses induced by impulsive shear flow from single cavitation microbubbles (SCBs). We used HEK293T cells with Piezo1 genetically knocked out (P1KO) or transiently transfected (P1TF).^[42]^ We treated the cells with integrin-binding Arginylglycylaspartic acid (RGD)-coated microbeads together with antibodies that block integrin ligation or with a P2X purinoreceptor inhibitor to dissect which of these previously-mentioned molecular players are involved. We found that the RGD beads were required to enhance the mechanical coupling and elicit a Ca^2+^ response without membrane poration. We established that the cellular mechanical sensing induced by microbubbles is mediated by increased integrin ligation, which leads to release of extracellular ATP (eATP) and subsequent activation of P2X channels, resulting in Ca^2+^ entry and downstream change in cell spreading. We anticipate that the insights from this study will enable development of new strategies for better use of therapeutic ultrasound.

## 2. Results

### 2.1. Impulsive Shear Flow Induced by Single Cavitation Bubbles

Focused laser pulses were used to generate SCBs that stimulated multiple isolated cells simultaneously, increasing the throughput of the experiment. The glass bottom of the petri dish was coated with a thin layer of gold to promote SCB nucleation, and with fibronectin (FN) to promote cell adhesion. This experimental setup allowed us to assess the response of individual cells located at various distances from a single microbubble within the same flow field (**Figure 1***A–C*). Normalized standoff distances (γ) were defined as γ = S_d_/R_max_, where S_d_ is the cell-SCB distance, and R_max_ is the maximum bubble radius. To assess the role of Piezo1, a mixture of P1KO and P1TF cells was seeded; these cells were distinguished by green fluorescence from GFP co-expressed with Piezo1 in P1TF cells.^[42]^ For more details on the experimental setup, see Experimental Section and *SI Appendix*, Figure S1.

**Figure 1.**
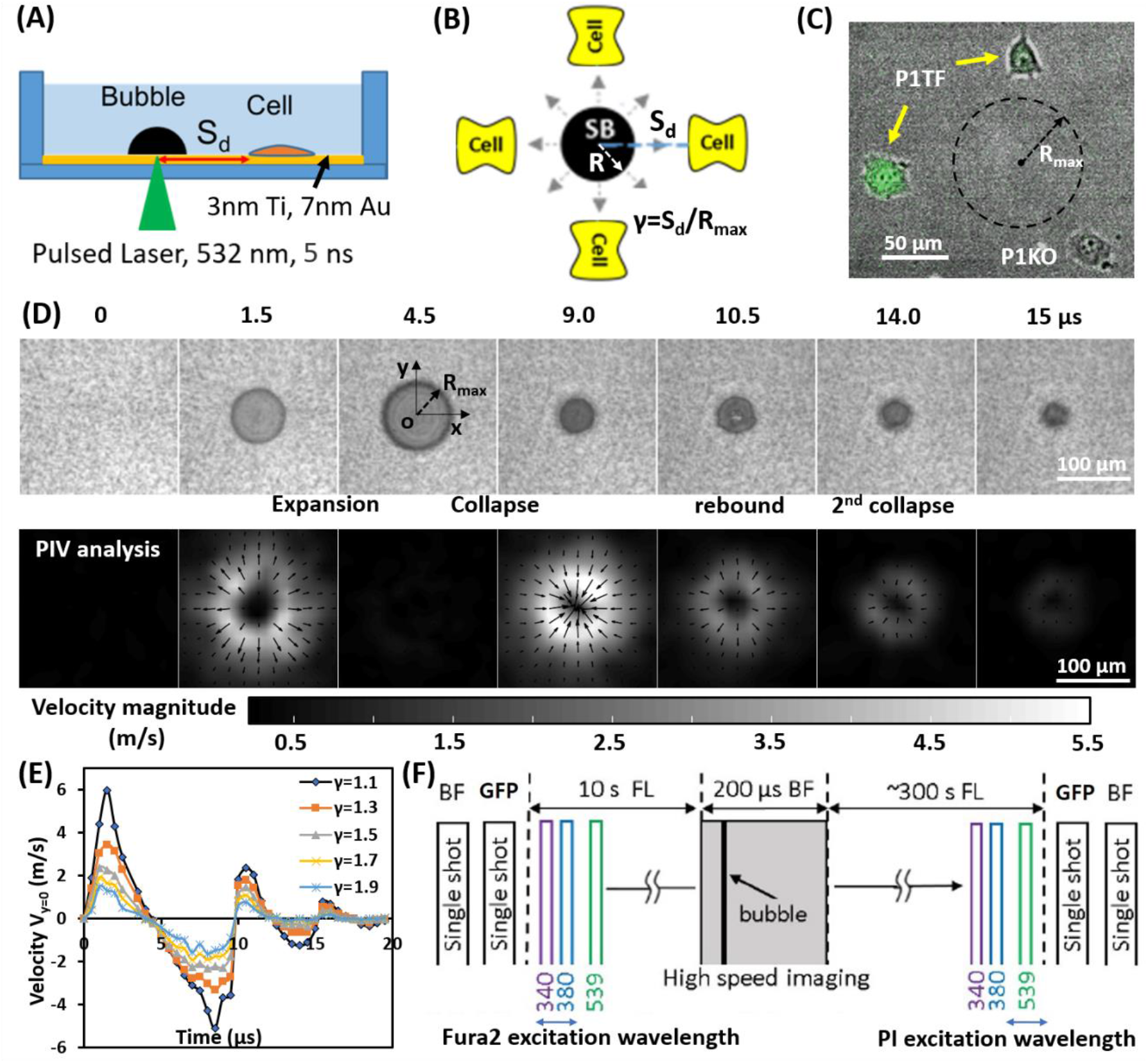
Experimental design. (A) A schematic of experimental setup. A laser-induced single cavitation bubble (SCB) stimulates nearby cells in a glass-bottom dish, which is coated with 3 nm Ti and 7 nm Au to enhance laser absorption while allowing optical transmission for microscopy. (B) Top view of experimental setup. S_d_, standoff distance; γ, normalized standoff distance; R_max_, maximum bubble radius. (C) Superimposed bright-field and fluorescence image of three cells with different S_d_ to a SCB with R_max_ indicated by black dashed line. A mixture of stable Piezo1-knockout cells (P1KO) and cells transiently transfected with GFP-Piezo1 (P1TF) was used. (D) High-speed images of bubble dynamics (upper panel) and velocity field from particle image velocimetry (PIV) (lower panel). (E) Time evolution of the SCB-generated impulsive radial flow velocity along a horizontal axis (y=0) through the bubble center. Outward velocity is positive, inward velocity is negative. The velocity amplitude decreases with γ. (F) Recording sequence: bright-field imaging of cell morphology, GFP imaging of Piezo1 expression, fluorescence imaging to simultaneously monitor intracellular Ca^2+^ transients (340- and 380-nm excitation) and membrane poration (539-nm excitation), and high-speed imaging of SCB dynamics.

In each set of measurements, a focused laser pulse nucleates a transient SCB that expands to its maximum radius in ~4.5 μs then collapses, followed by a second expansion and collapse with greatly reduced strength within 20 μs (Figure 1*D* and *E*). Particle image velocimetry (PIV) measurements revealed that the velocity of the SCB-generated impulsive radial flow decreases with increasing standoff distance, ranging from a maximum of ~6 m s^−1^ at γ = 1.1 to ~1.6 m s^−1^ at γ = 1.9 (Figure 1*E*). The radial flow imposes transient shear stress on adherent cells nearby (γ = 1–2) (*SI Appendix*, Figure S2), as shown previously.^[43]^ Laser-induced SCBs thus allow tuning the magnitude of shear flow applied to cells by adjusting the standoff distance.^[30, 44]^

Ca^2+^ influx following cell exposure to cavitation bubbles may occur either through ion channels or through poration of the plasma membrane.^[30]^ To distinguish these pathways, we used time-elapsed fluorescence imaging to concurrently monitor SCB-induced intracellular Ca^2+^ transients (with ratiometric imaging of Fura-2 excited at 340 and 380 nm) and membrane poration (with excitation of propidium iodide (PI) at 539 nm). The baseline fluorescence was recorded for 10 s, followed by high-speed imaging of bubble dynamics, then a ~300 s sequence of fluorescence imaging of the Ca^2+^ response and PI uptake in a cell (Figure 1*F*). Cell morphology was characterized before and after bubble treatment with single-shot bright-field images. Cell injury was assessed by PI uptake and cell morphology change.

### 2.2. SCB-Elicited Ca^2+^ Responses at Various Distances

Figure 2 presents time series images of intracellular Ca^2+^ response and PI uptake in individual P1KO and P1TF cells exposed to SCBs at a standoff distance of γ = 1.0–1.5. Bright-field images of cell morphology before and after SCB treatment are also shown. The average Ca^2+^ response was quantified as (F-F_0_)/F_0_, where F=I_340_/I_380_, and F_0_ is the value before bubble treatment. For both P1KO and P1TF cells, distinct Ca^2+^ responses were observed depending on the cell injury. Fast Ca^2+^ responses with a short rise time and a large amplitude change were accompanied by PI uptake associated with membrane poration (Figure 2*A*, *C* and *E*, red and blue traces). Slower and milder Ca^2+^ responses were accompanied by negligible PI uptake (little or no membrane poration) (Figure 2 *B*, *D* and *E*, orange and cyan traces). The Ca^2+^ response amplitude increased with PI uptake (*SI Appendix*, Figure S3), consistent with our previous findings when using tandem bubbles to stimulate HeLa cells.^[30]^

**Figure 2.**
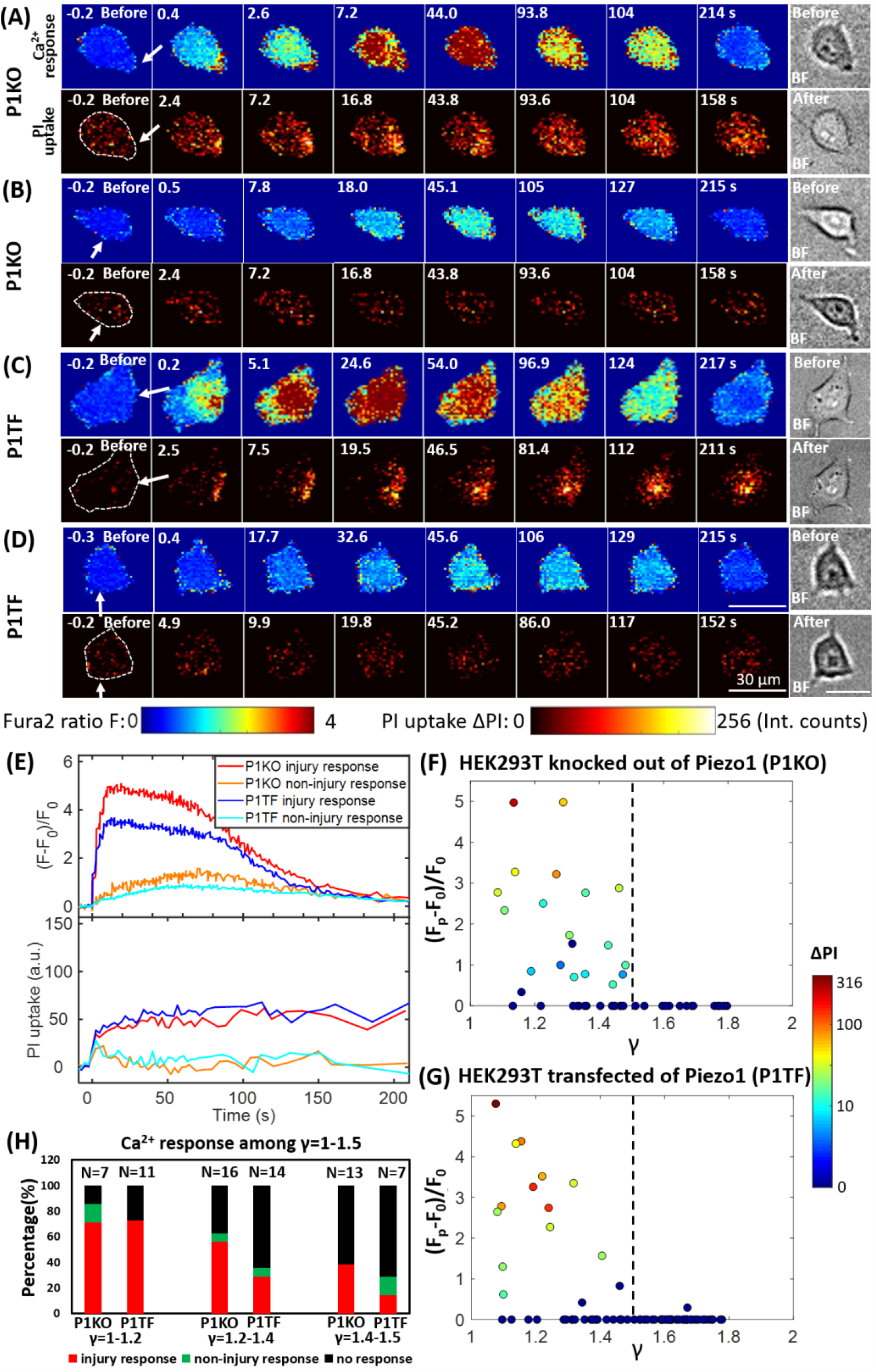
Intracellular Ca^2+^ response and PI uptake in SCB-stimulated cells. (A-D) Time series showing intracellular Ca^2+^ response (top rows) and PI uptake (bottom rows) in representative individual cells with membrane poration (A, C) or without membrane poration (B, D), for Piezo1 knockout cells (P1KO) (A-B) and cells transiently expressing Piezo1 (P1TF) (C-D). Right: Single-shot bright-field images of cells before and after treatment. The white arrow indicates the direction of radial expansion of the SCB at the location closest to the cell. Left: The cell contour before treatment is outlined with white dashed lines in the first PI image. Cells were exposed to SCB at t=0. Scale bars, 30 μm. (E) Temporal profiles of averaged intracellular Ca^2+^ response and PI uptake inside each cell shown in A–D. (F) and (G) Peak intracellular Ca^2+^ response (circles) versus standoff distance for P1KO and P1TF cells. PI uptake values are color-coded. Almost all elicited Ca^2+^ responses occurred at *γ* = 1–1.5. (H) Overall Ca^2+^ response probability at *γ* = 1–1.5, including cells with an injury response (red), non-injury response (green), and no response/no injury (black) at different ranges of *γ* (1–1.2, 1.2–1.4, and 1.4–1.5). N indicates the number of cells in each group.

Since the shear flow strength decreases with γ, we examined the effect of γ on the peak Ca^2+^ response, (F_p_-F_0_)/F_0_, and on PI uptake. Most of cells exhibiting a Ca^2+^ response were located within γ = 1.0–1.5 of the SCB, for both P1KO and P1TF cells (Figure 2 *F* and *G*). Ca^2+^ responses within the range of γ = 1.0–1.5 were further classified based as “injury response” (PI uptake), “non-injury response” (no PI uptake), and “no response”. The frequencies of these responses in P1KO and P1TF cells were similar: 53% and 41% for injury response, 5% and 6% for non-injury response, and 42% and 53% for no response for P1KO and P1TF cells, respectively (Figure 2*H* and *SI Appendix*, Figure S3). As expected, the fraction of no-response cells increased at larger standoff distances where shear stress was smaller (Figure 2*H*). Within γ = 1–1.5, there were more injury than non-injury responses. Since there was no significant difference in Ca^2+^ response between Piezo1 knockout and Piezo1-expressing cells, Piezo1 does not contribute to Ca^2+^ responses elicited by SCB-induced impulsive shear flow.

### 2.3. Enhancing Ca^2+^ Responses Without Cell Membrane Poration with RGD Beads

An alternative approach to elicit a Ca^2+^ response is to enhance the mechanical load applied to cells at standoff distances at which no membrane poration occurs (γ = 1.5–1.8). Informed by our previous study, we explored the effects of attaching 6-μm diameter RGD-coated polystyrene microbeads to the apical surface of the cells.^[30]^

**Figure 3***A* depicts a high-speed imaging sequence of SCB-cell interaction and the resultant displacement of RGD microbeads attached to the cell. Enlarged images (Figure 3*B*) show that the bead closest to the bubble was displaced significantly from its original position. The bead moved away from its original position during bubble expansion, and back to its original position following bubble collapse, over a period of ~22 μs. The corresponding time-elapse images of Ca^2+^ response and PI uptake (Figure 3*C*) indicate a non-injury Ca^2+^ response.

**Figure 3.**
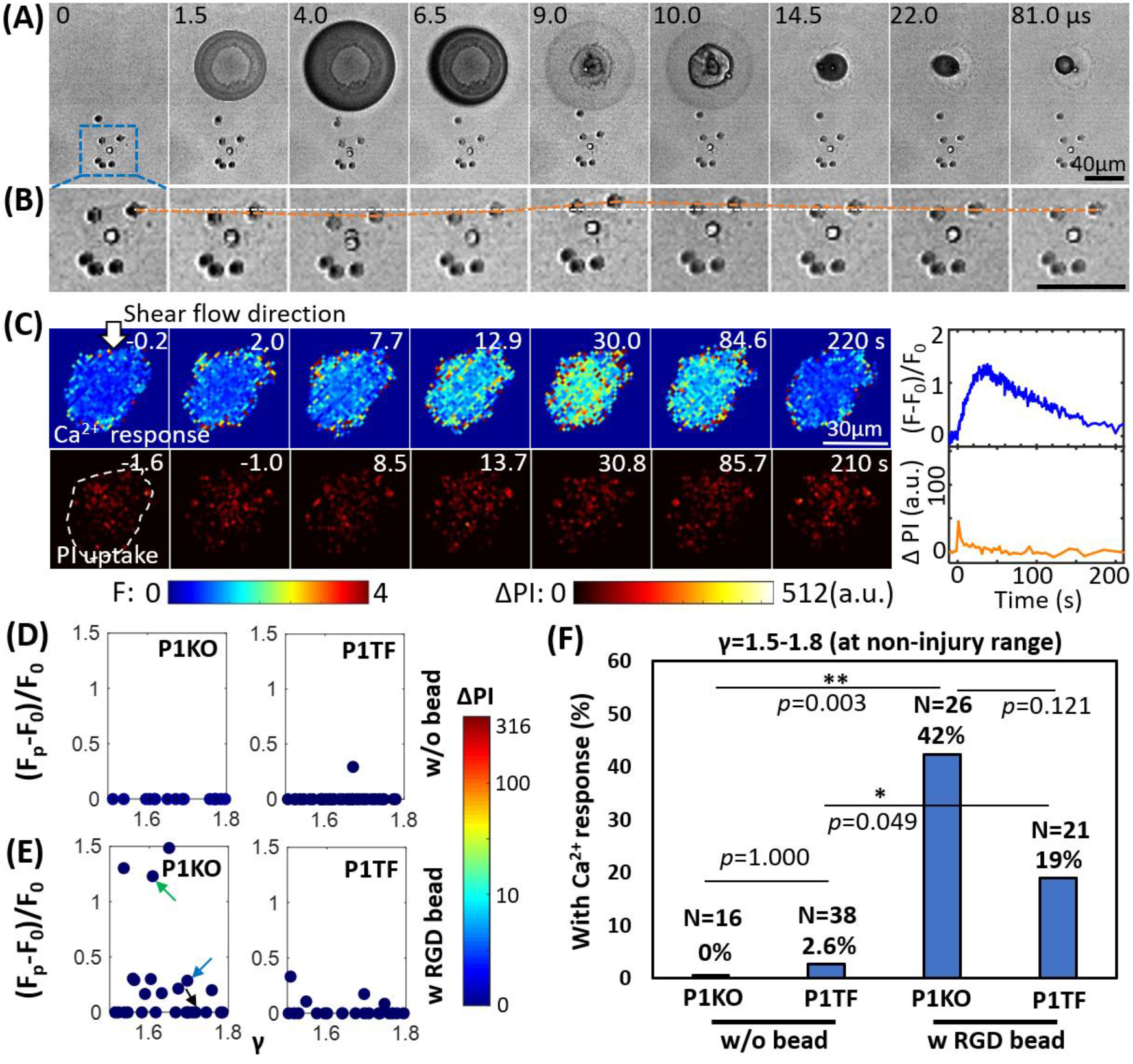
RGD beads enhance the Ca^2+^ response in the non-injury range of SCB treatment (γ = 1.5–1.8). (A) Image sequences from high-speed imaging showing an SCB-cell interaction with RGD beads attached to integrins on the cell apical surface. (B) Enlarged image sequence corresponding to the dashed box in A, showing displacement of the bead closest to the center of the SCB (upper right of the cell). The white line indicates the bead’s original position; the orange line indicates its position at subsequent time points. (C) *Left*: Time-elapsed image sequences of Ca^2+^ response without PI uptake in a cell subjected to SCB treatment with RGD beads. *Right*: Time traces of the normalized Ca^2+^ response and PI uptake. (D) and (E) Dependency of normalized peak Ca^2+^ response on γ with the amount of PI uptake color-coded, for P1KO and P1TF cells with and without RGD beads. γ = 1.5–1.8. No cells show PI uptake, indicating no membrane poration. (F) Percentage of P1KO and P1TF cells exhibiting a Ca^2+^ response, under SCB treatment within γ = 1.5–1.8, with or without RGD beads. \**p*<0.05, ** *p*<0.01, Fisher’s exact test. N is the number of cells in each treatment group. The *p* value of fisher exact probability test between the four groups is less than 0.0001.

Attachment of RGD beads enhanced the probability of eliciting a Ca^2+^ response at a normalized cell-bubble distance of γ = 1.5–-1.8 (Figure 3*E*). Without RGD beads, almost no cells showed a Ca^2+^ response at this distance (Figure 3*D*). Notably, no PI uptake was observed in the RGD bead-treated cells—all Ca^2+^ responses were non-injury responses, for both P1KO and P1TF cells. The fraction of cells showing a Ca^2+^ response increased significantly, from 0% without beads to 42% with beads (p<0.01) for P1KO, and from 2.6% to 19% (p<0.05) for P1TF (Figure 3*F*). No statistically significant difference was found between P1KO and P1TF cells (*p*>0.05).

Attaching integrin-binding microbeads is thus an efficient approach to eliciting Ca^2+^ responses with impulsive shear flow without cell injury. Again, no significant difference in Ca^2+^ response between Piezo1 knockout and Piezo-expressing cells was observed, indicating Piezo1 is not involved in the Ca^2+^ response.

### 2.4. Increased Integrin Ligation to Substrate ECM Initiates Ca^2+^ Signaling from the Cell Periphery

To discern the processes that initiated the Ca^2+^ response to SCB with RGD beads, we examined spatiotemporal changes in Ca^2+^ signaling within P1KO cells at distances causing a non-injury response (γ = 1.5–1.8). **Figure 4***A* shows a contour map of Ca^2+^ signaling amplitude in a representative P1KO cell. Interestingly, the Ca^2+^ signaling appeared to propagate from the cell periphery toward the center, consistent with a previous study.^[45]^ To confirm this observation, we segmented the cell into three radial regions of interest (ROI) and measured the Ca^2+^ response, (F-F_0_)/F_0_, in each ROI versus time (Figure 4*B*, *left*). The outermost ROI reached 50% of its maximum Ca^2+^ response earlier than the center ROI (~7.2 s versus ~12.5 s), with the Ca^2+^ wave propagating from the edge to the center of the cell at ~2.3 μm s^−1^ (Figure 4*B*, *right*).

**Figure 4.**
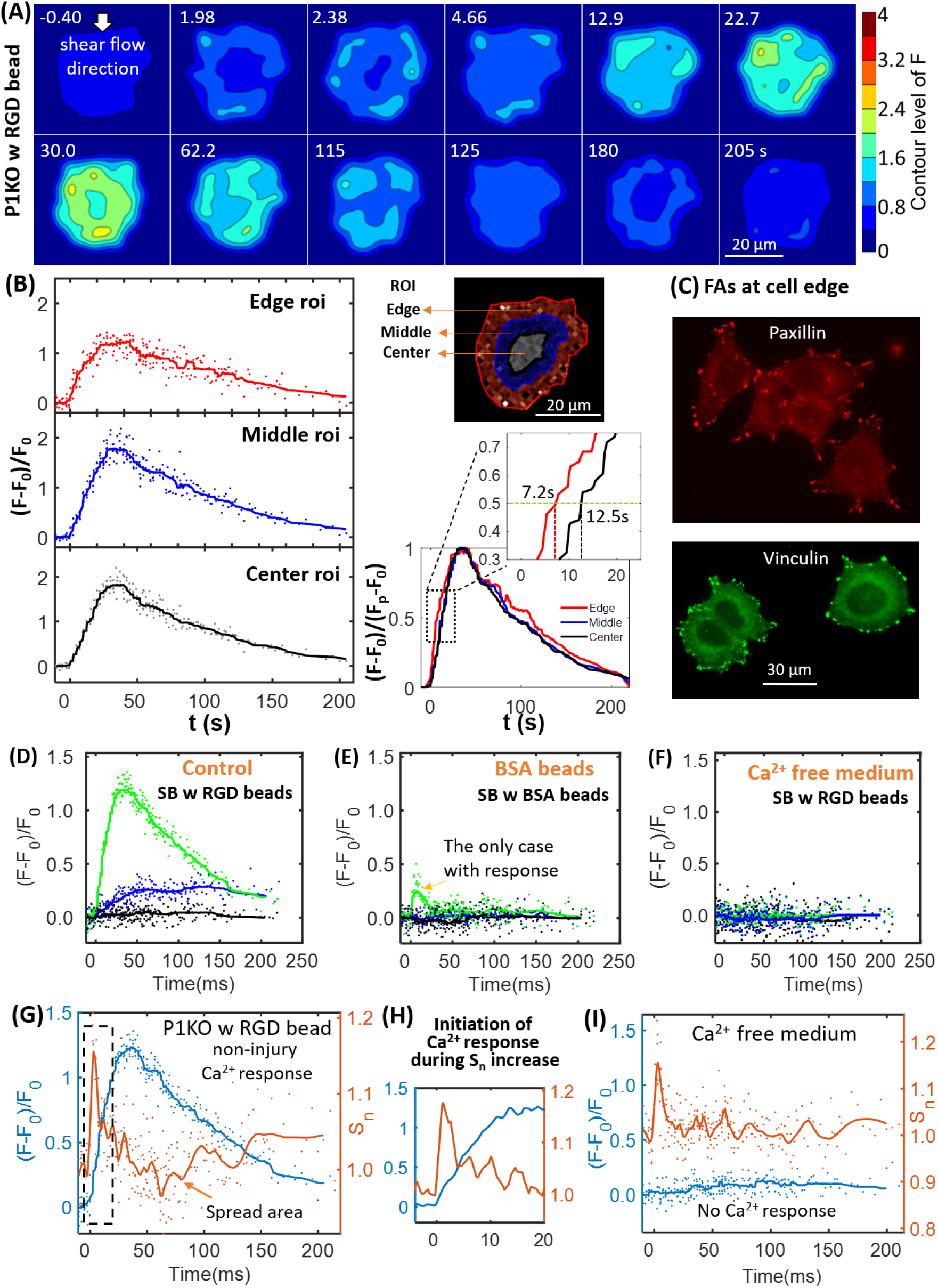
Initiation of the Ca^2+^ wave and localization of the focal adhesion molecules paxillin and vinculin at the cell periphery, and Ca^2+^ influx during the increase in cell spread area. (A) Surface contour of F=I_340_/I_380_ (ratiometric imaging of Ca^2+^ signaling amplitude) for a P1KO cell exhibiting a non-injury Ca^2+^ response, with RGD beads (the same cell as in Figure 3*C*). (B) Small region analysis (edge, middle and center ROI) of the Ca^2+^ response for the cell in panel A. *Left*: Time traces of (F-F_0_)/F_0_ and median averaging. *Top right*: ROIs labels. *Bottom right*: A normalized version of the median averaging line traces is shown in the dashed box around (F-F_0_)/(F_p_-F_0_)=0.5 (horizontal dashed line). (C) Fluorescence images of focal adhesion proteins paxillin and vinculin in P1KO cells after 3 h of spreading on a gold- and FN-coated dish. (D-F) Time traces of (F-F_0_)/F_0_ for SCB-treated P1KO cells with (i) RGD beads, (ii) BSA beads, and (iii) RGD beads in Ca^2+^-free medium. (G) Time traces of Ca^2+^ response and normalized change in cell spread area (Sn) for a SCB- and RGD-bead treated P1KO cell exhibiting a non-injury Ca^2+^ response. Solid lines indicate median averages (see Experimental Section). (H) Enlarged image of the dashed box in panel G showing that the Ca^2+^ response is initiated during the increase in Sn. (I) Time traces of (F-F_0_)/F_0_ and Sn for a no-response case in Ca^2+^-free medium, with a similar transient increase in Sn as seen in panel H, indicating that Ca^2+^ entry is necessary for RGD bead-enhanced response. All cases shown are cells stimulated with SCB at γ = 1.5–1.8.

Since cells detect extracellular mechanical stimuli through integrins and focal adhesions,^[36, 46]^ we investigated whether the Ca^2+^ response is integrin-specific. Paxillin and vinculin are adaptor proteins in focal adhesions that connect transmembrane integrins with the actin cytoskeleton.^[47]^ Based on immunofluorescence staining before bubble treatment, both paxillin and vinculin were localized mainly at the cell periphery (Figure 4*C*). We examined whether integrin binding is required on the cell apical surface by comparing Ca^2+^ responses of P1KO cells within γ = 1.5–1.8 exposed to beads coated with either RGD or BSA (Figure 4 *D* and *E*). BSA-coated beads non-specifically bind cell membranes but experience the same drag as RGD-coated beads. The Ca^2+^ response was significantly reduced in cells when using BSA beads, indicating that integrin-specific bead attachment was vital to the Ca^2+^ response. No Ca^2+^ response was observed when using Ca^2+^-free medium with RGD beads (Figure 4*F*), indicating that the increased cytosolic Ca^2+^ was dependent upon extracellular Ca^2+^ entry.

Increased ligation of integrins by the substrate ECM has been associated with shear-induced mechanotransduction.^[33–36]^ We hypothesized that increased integrin ligation may be responsible for initiating the Ca^2+^ response in our study. Therefore, we examined cell spreading as an indicator for integrin-ECM interactions following SCB treatment. Normalized cell spread area was quantified as S_n_ = S/S_0_, where S_0_ is the area before SCB treatment. We observed a transient increase in Sn immediately after SCB treatment, from 2.0 s to 2.4 s (Figure 4 *A* *and* *G*). A Ca^2+^ response was initiated concurrently with this increase in S_n_ (Figure 4 *G* and *H*). In contrast, in Ca^2+^-free medium, a similar increase in S_n_ occurred but was not associated with a cytosolic increase in Ca^2+^ (Figure 4*I*), suggesting that the integrin ligation-induced Ca^2+^ response requires extracellular Ca^2+^ entry.

### 2.5. SCB-Induced Ca^2+^ Response Drives Transient Reduction in Cell Spread Area

After the transient increase in cell spreading immediately following SCB treatment, the spreading area decreased and then recovered to a plateau at ~150 s (Figure 4*G*). We examined this process in detail for three P1KO cells at γ = 1.5–1.8 with RGD beads before extending the analysis. The three cells showed three different Ca^2+^ responses (**Figure 5***A*–*C*). In the two cases with non-injury responses, the minimum spreading area (Sn)min was reached soon after or at nearly the same time as the Ca^2+^ peak (Figure 5 *A* and *B*). The reduction in cell spread area,1-(S_n_)_min_, was greater for the cell showing the larger Ca^2+^ response (~10% in Figure 5*A* vs. ~7.5% in Figure 5*B*), and was <2% when there was no Ca^2+^ response (Figure 5*C*). To reinforce this observation, we analyzed reduction in area for additional cells under the same conditions. Figure 5*D* and *E* summarize the dependence of area reduction on Ca^2+^ response and standoff distance, respectively. The reduction in area was <5% for no-response cases, but increased with increasing Ca^2+^ response, reaching a maximum of 11% (Figure 5*D*). No clear dependence on standoff distance was observed for the no-response cases (Figure 5*E*, black). In contrast, for non-injury response cases, the reduction in area decreased with increasing γ (Figure 5*E*, blue). Together, the results suggest that the reduction in cell spread area is a downstream effect of the Ca^2+^ response.

**Figure 5.**
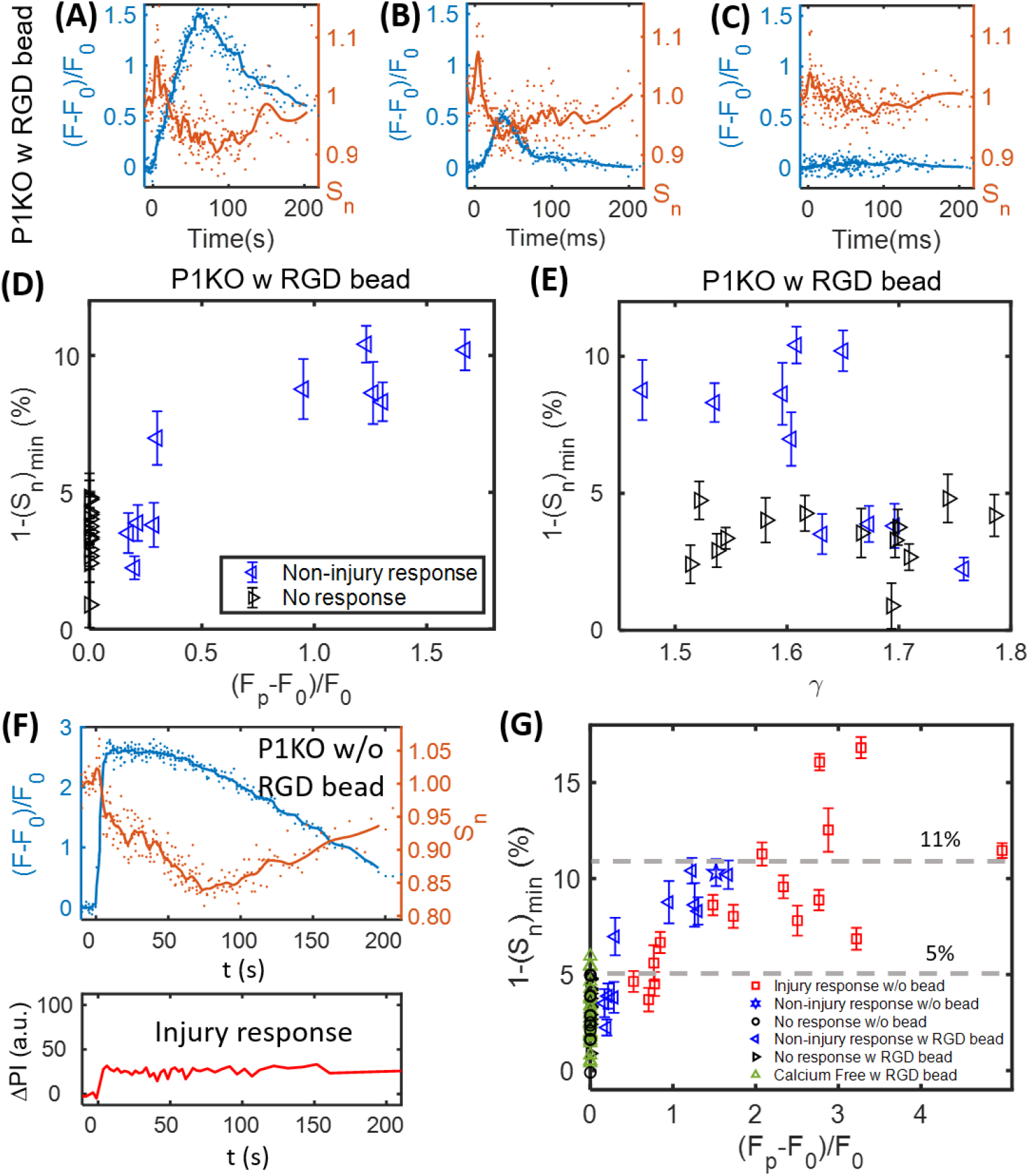
The SCB-induced Ca^2+^ response drives a transient reduction in cell spread area. (A-C) Change in Ca^2+^ response (blue traces) and normalized cell spread area (orange traces) versus time for three P1KO cells with RGD beads in the non-injury range (γ = 1.5–1.8). The peak Ca^2+^ response decreases from A to C. (D) Correlation of the peak Ca^2+^ response amplitude with the maximum reduction in cell spread area, 1-(S_n_)_min_, for each P1KO cell. (E) Distribution of 1-(S_n_)_min_ versus γ for the data shown in panel D. (F) Example of an injury Ca^2+^ response from SCB treatment of P1KO cells without RGD beads in the sub-lethal range of γ = 1–1.5, showing the change in cell spread area, Ca^2+^ response, and PI uptake over time. (G) The maximum reduction in cell spread area with Ca^2+^ response amplitude for P1KO cells with and without RGD beads. The maximum reduction in cell spread area depends on the amplitude of the Ca^2+^ response. The error bars in panel D, E and G depict the SEM.

We also examined whether the reduction of cell spread area occurs in other Ca^2+^ response cases. In a P1KO cell without RGD beads that exhibited an injury Ca^2+^ response, the Ca^2+^ signal rose sharply upon SCB treatment at 0 s (Figure 5*F*), a response that is distinct from the non-injury responses (Figure 5*A* and *B*). However, there was little transient increase in cell spreading area. This result agreed with our previous finding that extracellular Ca^2+^ influx occurs through pores in the plasma membrane in injury response cases.^[30]^ Despite the different route of Ca^2+^ entry, in both cases we observed a reduction and recovery of cell spread area following the Ca^2+^ response.

To assess whether Ca^2+^ influx and integrin-binding RGD beads are necessary to induce the reduction in cell spreading, we extended the analysis to cases of P1KO cells with Ca^2+^-free medium or without RGD beads (Figure 5*G*). There was no Ca^2+^ response in Ca^2+^-free medium, as observed previously. In all cases with no Ca^2+^ response, the reduction in cell area was <6% and was often comparable to the error from measurement (1.8% - 4.3% standard deviation (SD), and 0.4%-0.9% standard error of the mean (SEM)), indicated no significant change. In contrast, for cells exhibiting a Ca^2+^ response, the reduction in cell area increased with the Ca^2+^ response amplitude and reached higher levels (>15%).

This analysis suggested that the transient decrease in cell spread area is a downstream effect of the Ca^2+^ response; i.e. cell spreading is a result of ‘inside-out’ signaling from regulation of cell adhesion by increased intracellular Ca^2+^. Additional analysis of our previous data also showed transient reduction in cell spread area for tandem bubble-treated HeLa cells with or without RGD beads, which had a Ca^2+^ response (*SI Appendix*, Figure S4).^[30]^ This reduction in area may be caused by myosin-II-dependent cell contraction and/or intracellular Ca^2+^-dependent calpain activation that leads to focal adhesion disassembly.^[48, 49]^

### 2.6. RGD Bead-Enhanced Ca^2+^ Response Requires New Integrin Ligation and Subsequent Extracellular ATP Release

We investigated whether new ligation of integrins at the cell-ECM substrate interface is necessary to initiate the Ca^2+^ response. After P1KO cells had spread on FN-coated dishes, the FN-specific antibody 16G3 was added to block any new integrin α_v_β_3_ and α_5_β_1_ binding sites. In this way, the cells could only maintain previously established integrin-ECM connections during SCB treatment. A non-blocking FN-specific antibody, 13G12, was used as a control. Treatment with 16G3 completely suppressed the RGD bead-enhanced Ca^2+^ response, while 13G12 had no significant effect (*p*>0.05) on the Ca^2+^ response (**Figure 6**A and B, and **Figure 7**A).

**Figure 6.**
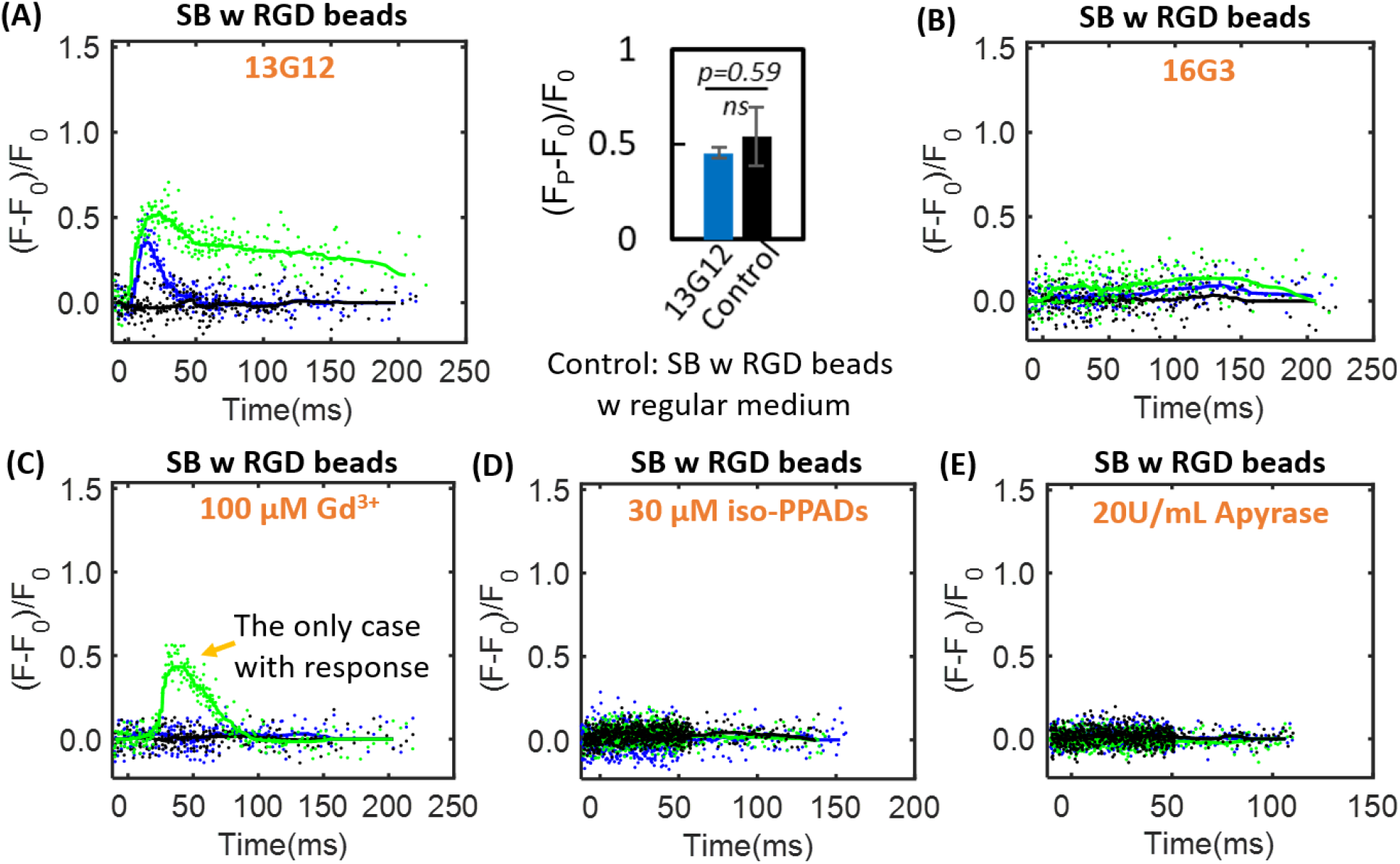
Representative time traces of Ca^2+^ response for the mechanistic study of RGD beads enhanced Ca^2+^ signaling. (A) and (B) HEK293T P1KO cells with RGD beads and treated with fibronectin antibody 13G12 (w/o blocking integrin ligation, 20 μg mL^−1^) and 16G3 (blocking integrin-ligation, 20 μg mL^−1^) when subjected to SB induced flow, respectively. The bar graph in (A) shows the mean value of the amplitude of the normalized Ca^2+^ response (F_p_-F_0_)/F_0_ with error bar indicating the SEM, demonstrating no significant difference between 13G12 treated group and the control group with regular cell medium (two-tailed t-test). (C) SB treated P1KO cells attached with RGD beads with 100 μM Gd^3+^ in extracellular medium. The response case is the only cell with Ca^2+^ response among the whole treated population (N=15). (D) and (E) SB treated P1KO cells that are attached with RGD beads and incubated with 30 μM iso-PPADs to block P2X channels and 20 U mL^−1^ apyrase to deplete extracellular ATP, respectively.

**Figure 7.**
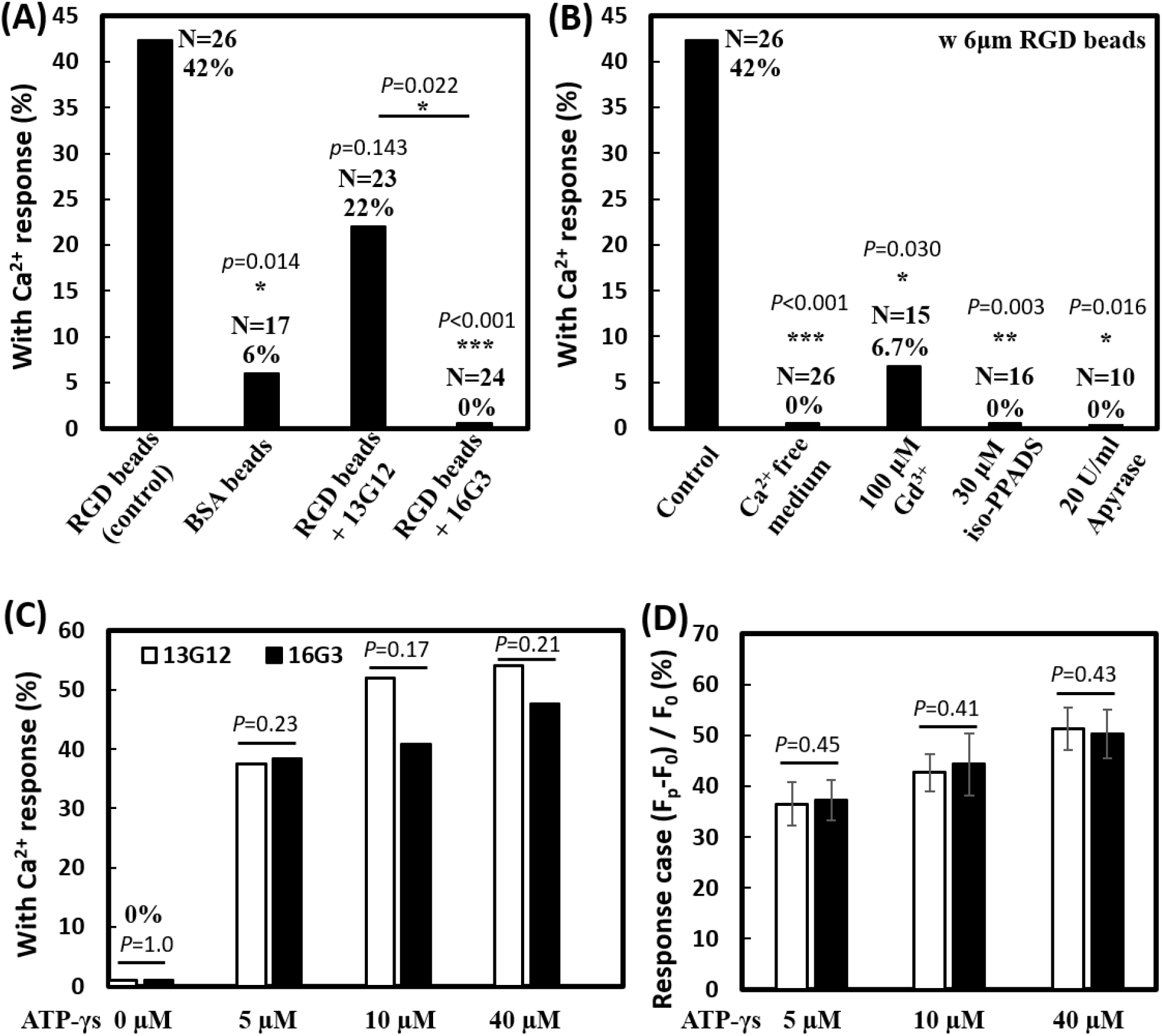
RGD bead enhancement of the non-injury Ca^2+^ response requires new integrin ligation and subsequent eATP release. (A) Percentage of non-injury Ca^2+^ response for SCB-treated P1KO cells at γ = 1.5-1.8 with (i) RGD beads; (ii) BSA beads; (iii) RGD beads plus non-integrin blocking anti-FN antibody 13G12 (20 μg mL^−1^); and (iv) RGD beads plus integrin-blocking antibody 16G3 (20 μg mL^−1^). (B) Percentage of non-injury Ca^2+^ response in P1KO cells with RGD beads with: (i) no additional treatment (control); (ii) Ca^2+^-free medium; (iii) 100 μM Gd^3+^ (GdCl3); (iv) 30 μM iso-PPADs (P2X channel inhibitor); and (v) apyrase (20 U mL^−1^) (depleting eATP). (C-D) P1KO cells were treated with anti-FN antibody 13G12 (white bars) or 16G3 (black bars) (20 μg mL^−1^) after spreading on FN-coated glass, then treated with different concentrations of ATP-γ-S in 1x PBS with Ca^2+^. The resulting percentage of Ca^2+^ response is shown in panel C); the average normalized Ca^2+^ response is shown in panel D (error bar indicates SEM). The total number of cells tested in panel C was N = 24, 24, 25, and 24 for 13G12 at 0, 5, 10, and 40 μM ATP-γ-S, respectively; N = 26, 26, 22, and 21 for 16G3 at 0, 5, 10, and 40 μM ATP-γ-S, respectively. The number of cells tested in panel D was N = 9, 13, and 13 for 13G12; N = 10, 9, and 10 for 16G3 at 5, 10, and 40 μM ATP-γ-S. \**p*<0.05, *** *p*<0.001. *P*-values were calculated with two-tailed Fisher’s exact test in A-C, and one-tailed student’s t test in D. The *p* value of fisher exact probability test is less than 0.001 between the four groups in (A) and less than 0.0001 between the five groups in (B).

We examined whether the integrin-dependent, non-injury Ca^2+^ response resulted from extracellular Ca^2+^ influx through ion channels on the cell membrane. 42% of P1KO cells showed a non-injury Ca^2+^ response with RGD beads (Figure 7B). Extracellular Ca^2+^ influx is essential in these responses, as shown by using Ca^2+^-free medium (Figure 5*G* and Figure 7*B*). Gd^3+^ was shown previously to block both mechanosensitive ion channels and the ligand-gated P2X channel;^[50]^ in our study, addition of 100 μM Gd^3+^ also significantly suppressed the Ca^2+^ response (Figure 6C and Figure 7*B*). Since our results suggest that the mechanosensitive ion channel Piezo1 is not critically involved under the impulsive shear flow (Figure 3*F* and Table S1), we hypothesized that the P2X channel is vital for the extracellular Ca^2+^ influx. P2X channels have been shown to be responsible for shear stress-generated Ca^2+^ waves that propagate from the edge to the center of rat atrial myocytes.^[45]^ This hypothesis was supported by adding the P2X channel-specific blocker pyridoxalphosphate-6-azophenyl-2’,5’-disulfonic acid (iso-PPADS, 30 μM), which eliminated the Ca^2+^ response (Figure 6D and Figure 7*B*).

P2X is an ATP-gated channel. We depleted eATP with 20 U mL^−1^ apyrase (an ATP-diphosphatase) and found that the Ca^2+^ response was completely suppressed (Figure 6E and Figure 7*B*), reinforcing the hypothesis that P2X activation mediates Ca^2+^ influx following SCB treatment. We next established the causality between integrin ligation and eATP release, as both are required for the non-injury Ca^2+^ response. If integrin ligation is downstream of eATP release, the Ca^2+^ response following eATP stimulation should be affected by blocking integrin-ECM binding sites with 16G3 antibody. P1KO cells were allowed to spread on FN-coated coverslips before treatment with 16G3 or non-blocking 13G12 antibodies. The cells were then stimulated with a non-hydrolyzable ATP analog, ATP-γ-S, at 5–40 μM. We observed a dose-dependent increase in the Ca^2+^ response with ATP-γ-S treatment (Figure 7 *C* and *D*). However, there was no significant difference in the Ca^2+^ response between 16G3- and 13G12-treated cells stimulated with ATP-γ-S, suggesting that integrin ligation is upstream of eATP release, consistent with previous studies.^[51–53]^

Altogether, our results suggest the following model for the SCB-elicited Ca^2+^ response (**Figure 8**): SCB-induced shear force is exerted on integrins on the cell apical surface by pulling on RGD-coated beads, causing new ligation of integrins by ECM proteins on the cell basal surface, which in turn triggers cellular release of eATP, which opens P2X ion channels, allowing Ca^2+^ influx that regulates a dynamic change in cell spreading.

**Figure 8.**
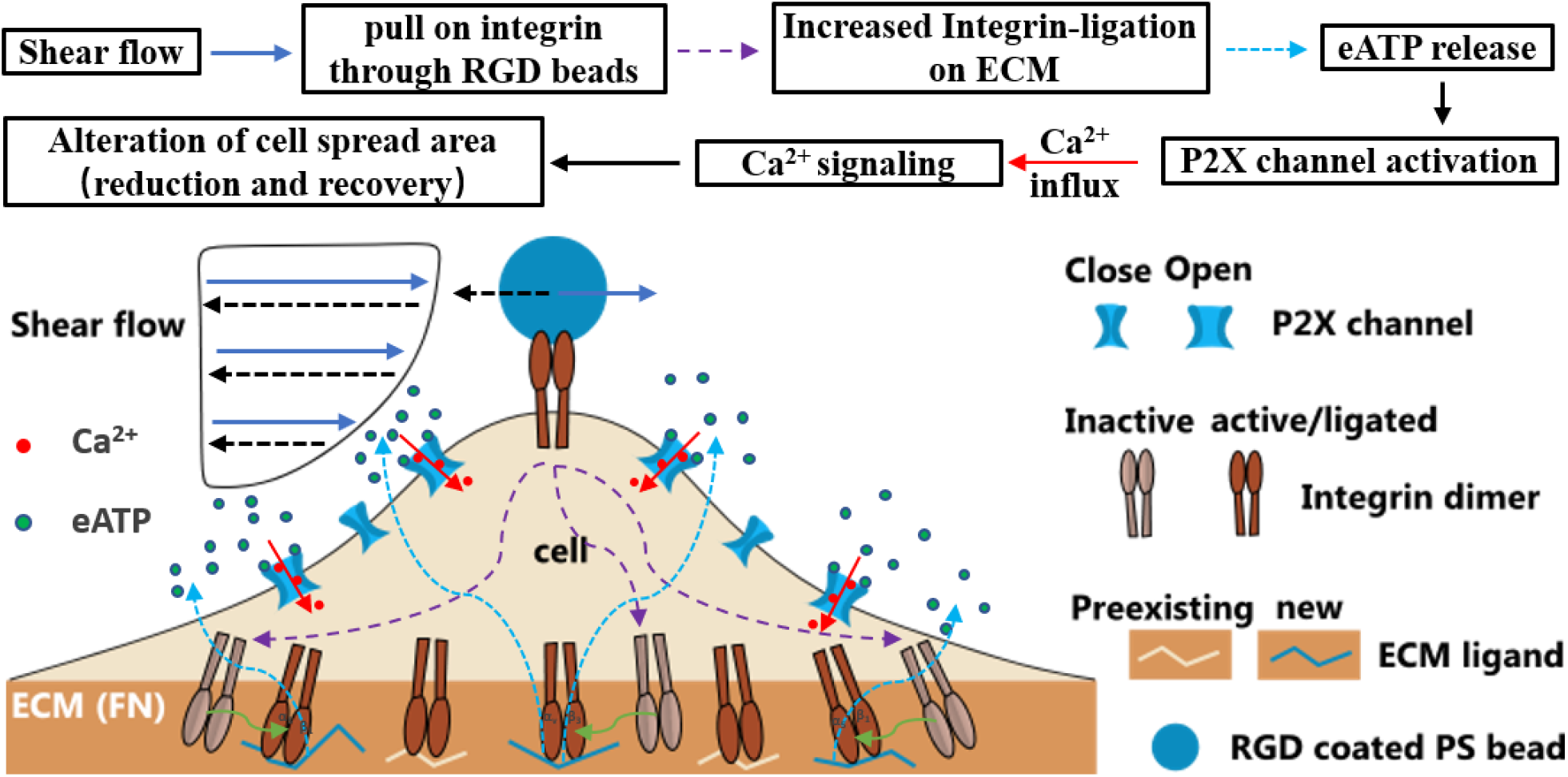
Schematic showing overall mechanotransduction process and key molecular players.

## 3. Discussion

In therapeutic ultrasound and shockwave applications, the shear force experienced by cells likely involves a high strain rate.^[13, 54]^ The effects of this impulsive shear force on cell mechanotransduction and Ca^2+^ signaling have not been previously examined. In this study we developed a system in which laser-generated SCBs produce impulsive shear flow on multiple nearby cells, allowing control of the distance between SCB and the cells, and used this system to dissect the key mechanisms underlying the elicited Ca^2+^ response.

In general, shear-induced mechanotransduction can be mediated by several different intracellular or extracellular structures, including mechanosensitive ion channels, purinergic receptors, and integrins, ECM.^[55–57]^ By attaching integrin-binding RGD microbeads to the cell apical surface, we reliably elicited Ca^2+^ signaling at SCB-cell distances that did not cause membrane poration. The mechanical loading applied to the integrin-bound beads led to increased integrin ligation to the substrate ECM, triggering eATP release that activated ATP-gated P2X channels, and resulting in Ca^2+^ influx and downstream signaling. This approach is also effective under impulsive jetting flow from tandem bubbles,^[30]^ where a similar integrin ligation mechanism is suggested (Figure S4 A-B). The involvement of integrin ligation by ECM and eATP release due to high strain rate loading has not been reported before. Moreover, under the same experimental conditions, BSA-coated beads did not induce Ca^2+^ signaling, suggesting that the signaling is not due to shear force transferred to the cell membrane or cell as a whole. Instead, mechanical transmission via integrins, likely linked to the cell basal surface via the cytoskeleton,^[37]^ appears to be necessary to transfer the stimulus to cell basal sites for activation of integrin-ligation and Ca^2+^ signaling.

The mechanosensitive ion channel Piezo1 has been previously implicated in Ca^2+^ responses in cells stimulated by quasi-static shear flow.^[31, 58]^ However, we observed no significant difference in Ca^2+^ response between Piezo1 knockout and Piezo1-expressing HEK293T cells, either with or without RGD beads under impulsive shear flow, indicating that Piezo1 is not involved under this shear loading regime. In general, Piezo1 enhances mechanotransduction at low strain rates and longer duration stimulations,^[17, 39, 40, 58, 59]^ but not at high strain rates due to impulsive shear flow from cavitation microbubbles, as shown here. The involvement of Piezo1 appears to be a key factor distinguishing cell mechanotransduction at different loading regimes.

Some elements of the mechanism identified in our study have been reported previously for ultrasound-induced mechanotransduction. For example, 2 MHz ultrasound alone did not evoke a Ca^2+^ response in HEK293T cells transfected with Piezo1, but did evoke a response when integrin-binding microbubbles were attached to the cell membrane.^[17]^ The importance of integrins in transducing mechanical forces produced by ultrasound has also been suggested by the effect of therapeutic low-intensity pulsed ultrasound (LIPUS) on cell motility via integrin-ECM adhesions.^[56]^ In our study, eATP release and purinergic signaling was important in Ca^2+^ response, and were also shown to be important in Ca^2+^ signaling in human mesenchymal stem cells stimulated using low-intensity ultrasound.^[60]^ Despite this scattered evidence, the molecular cascade identified in our study has not been reported in the context of ultrasound or cavitation microbubble induced mechanotransduction.

## 4. Conclusion

Our results establish that applying forces to integrins and activating ATP-gated P2X ion channels is an efficient approach to elicit a Ca^2+^ response in cavitation bubble-generated impulsive shear flow without membrane poration. This strategy does not require expression of exogenous mechanosensitive ion channels such as Piezo1 or MscL I92L,^[38]^ which are activated primarily by high-frequency ultrasound (tens of MHz), which has a limited penetration depth *in vivo*. Since expression of integrins and P2X is widespread in mammalian brain, muscle, and other tissues,^[55]^ ultrasound alone or mediated by cavitation could stimulate these tissues through a similar mechanism. In addition to neurotransmission and neuromodulation, P2X purinergic signaling is involved in cell proliferation, differentiation, motility, and apoptosis and in tissue regeneration and wound healing in muscle and other tissues.^[61, 62]^ By applying impulsive shear force with ultrasonic cavitation bubbles that deform the native ECM (thus pulling on integrins), we may be able to trigger the P2X purinergic signaling pathway and modulate cell functions in the brain and other tissues. Overall, these results provide insight into both the mechanism of cavitation bubble-mediated cell mechanotransduction and strategies for improving therapeutic ultrasound applications in tissue stimulation.

## 5. Experimental Section

### Petri Dish Surface Coating and SCB Setup

Glass-bottom petri dishes (35 mm, No. 1.5 coverslip, P35G-1.5-20-C; MatTek Corporation, Ashland, MA) were coated with a 3 nm layer of Ti and a 7nm layer of gold by electron beam evaporation. SCBs (maximum diameter, 90–110 μm) were produced in medium near cells by focusing a pulsed Neodymium-doped yttrium aluminum garnet (Nd:YAG) laser beam (*λ* = 532 nm, 5-ns pulse duration. Orion, New Wave Research) on the coated glass surface through a 20× objective (Fluar NA 0.75; Zeiss).

### Cell Culture, Piezo1 Transfection, and Cell Handling

The HEK293T-P1KO cell line (human embryonic kidney cells with Piezo1 genetically knocked out) and mouse Piezo1-pIRES-EGFP in pcDNA3.1 were obtained from Jorg Grandl’s lab at Duke Neurobiology, and have been described previously.^[42, 57, 63]^ P1KO cells were maintained in high-glucose DMEM with 10% heat-inactivated fetal bovine serum (FBS) and 1% penicillin/streptomycin at 37 °C and 5% CO_2_. At 48 h before SCB treatment and recording, P1KO cells were transiently transfected in a 6-well plate (the cells were seeded 8 h before transfection) in the presence of ruthenium red (10 μM) and mouse Piezo1 (3 μg) using Fugene6 (Promega, Madison, WI). 20–30% of cells showed positive GFP expression indicating successful Piezo1 transfection.

Two days after transfection, cells were seeded at low density in Au/Ti-coated dishes pre-wetted with 1× PBS and coated with FN (50 μg mL^−1^). Cells were incubated in DMEM at 37 °C for 3 h to allow cell adhesion and spreading. The culture medium was replaced with fura-2 AM (6 μM, F1221; Thermo Fisher Scientific) in Opti-MEM (11058-021; Thermo Fisher Scientific) and incubated at 37 °C in the dark for 30 min. Several (3–5) washes with 1× PBS were used to remove unloaded fura-2 AM before adding media in various experiments. PI was added at a final concentration of 100 μg mL^−1^ to monitor cell membrane poration.

In experiments with beads, cells were incubated at 37 °C for 30 min with 6-μm carboxyl-functionalized polystyrene beads activated with water-soluble carbodiimide, precoated with BSA (2.5% in PBS) or RGD-containing peptide (Peptite-2000; 100 μg mL^−1^ in PBS) in 1× PBS with Ca^2+^ (DPBS, Catalog No. 14040133, Gibco). Unattached beads were removed by washing with 1× PBS.

### Reagents and Treatment for Mechanistic Studies of RGD Bead-Enhanced Ca^2+^ Response

The following treatments were performed for mechanistic studies after fura-2 loading, RGD bead attachment, and washing with Ca^2+^-free 1× PBS: (1) Treatment with anti-FN antibodies followed protocols in the literature.^[36, 48]^ Cells were treated with 16G3 or 13G12 (20 μg mL^−1^) in 1× PBS with Ca^2+^ for 25 min at 37 °C before bubble treatment. (2) Cells were treated with Ca^2+^-free 1× PBS (DPBS, Catalog No.14190144, Gibco). (3) Gd^3+^ solution (100 μM GdCl_3_) in 1× PBS with Ca^2+^ was used. (4) For P2X channel inhibition, iso-PPAD (30 μM) in 1× PBS with Ca^2+^ was used. (5) To deplete extracellular ATP, cells were incubated with Apyrase (20 U mL^−1^) in 1× PBS with Ca^2+^ for 10 min before bubble treatment in the same medium.

### Determination of Whether Integrin Ligation is Upstream or Downstream of eATP Release

After 3 h of cell spreading on a gold-coated dish, P1KO cells were incubated with 13G12 or 16G3 (20 μg mL^−1^) in 1xPBS with Ca^2+^ for 25 min (37 °C). Cells were then treated with non-hydrolyzable ATP-γ-S (0(control), 5, 10, and 40 μM for separate dishes, respectively), and Ca^2+^ response was recorded with fura-2 ratiometric imaging.

### Image Recording and Analysis

Glass-bottom petri dishes with fura-2-loaded cells were fixed on the translation stage of an inverted microscope (Axio Observer Z1; Zeiss). Fluorescence excitation from a 75-W xenon lamp was controlled by a monochromator (DeltaRAM X; PTI) using shutters at defined time intervals, and at excitation wavelengths of 340 and 380 nm for fura-2 imaging, 539 nm for PI imaging, and 465 nm for GFP imaging. Since PI could also be excited at 340 and 380 nm, a custom-made narrow bandwidth filter (510 ± 40 nm) was used for detection of fura-2 emission at 510 nm to avoid overlap with PI emission at 610 nm. The intracellular fura-2 and PI images were recorded by using an sCMOS camera (EDGE 5.5 CL; PCO) at a 5:1 frame ratio, with exposure times of 50 ms and 100 ms, respectively. The image recording sequence for fura-2 was similar to our previous study;^[30]^ an interframe time (IFT) of 0.2 s for 1 min, then an IFT of 1, 2, and 5 s for 1, 1, and 2 min, respectively, for a total recording time of 300 s, during which one PI image was recorded every five frames of fura-2 images, under the control of μManager software (version1.4; Open Imaging).

SCB dynamics with or without bead displacement were captured by using a high-speed camera— either an Imacon 200 (DRS Hadland) or a Kirana K1 (Specialized Imaging, with SL-LUX640 laser for illumination)—at a recording rate of 2 million frames per second with 200-ns exposure time. The triggers for laser bubble generation and high-speed image acquisition were synchronized by a delay generator (565-8c; Berkeley Nucleonics Corporation). Fluorescence excitation, filter cubes, and image recording were automatically switched between fura-2 and PI with a switching interval ~0.7 s by μManager (Figure S1). Additional details of cell preparation, bubble generation, and imaging can be found in our previous studies.^[30, 64]^

Raw fura-2 images were corrected by background noise subtraction. The ratiometric value F=I_340_/I_380_ was calculated with MATLAB at each pixel before obtaining the averaged ratio within each cell or ROI from a manually segmented region of the cell with ImageJ. Time traces of the normalized ratio change (F-F_0_)/F_0_ for each cell were processed with a one-dimensional median filter in MATLAB to reduce the noise. F_0_ is the fluorescence ratio calculated by averaging the data before SCB treatment.

### Analysis of Cell Spread Area

Cell spread area was calculated from fura-2 images with MATLAB by thresholding at each combined image frame from 340 and 380 nm excitation. The emissions from these two wavelengths have opposite trends of fluorescence change; therefore, adding them for thresholding can reduce artifacts from fluorescence intensity change. The spread areas from the image frames before bubble treatment (S_0_) were averaged for normalization: S_n_ = S/S_0_. Time traces of S_n_ were processed with the one-dimensional median filter to reduce noise. The error analysis for cell spread area reduction is shown in *SI Appendix.*

### PIV Experiments and Analysis

Polystyrene beads (2 μm) in 1× PBS were used as tracers to map the flow field produced by SCBs. SCB dynamics and tracer bead movements were recorded by a Kirana camera with SL-LUX640 laser for illumination at 2 million frames per second and 100-ns exposure time. High-speed image sequences were analyzed offline using a commercial PIV software (DaVis 7.2; LaVision), then post-processed with MATLAB to obtain the time evolution of the flow velocity at different γ. For PIV software processing, the image field (370×370 μm) was divided into multiple interrogation windows of 20×20 μm each with 50% overlap, and multipass iterations and regional filters were applied to reduce the error in velocity field.^[29]^

### Immunostaining of Focal Adhesions

Cells were fixed with 4% formaldehyde for 10 min, permeabilized with 0.1% Triton X100 for 5 min, and blocked with 2% BSA for 1 h. Cells were then incubated with primary antibodies to vinculin (Sigma V9131) or paxillin (Abcam 32084) for 2 h, labeled with secondary antibodies (30 min), and imaged via fluorescence microscopy (Olympus IX83, UPlanSApo 60X/NA1.35 objective).

## Acknowledgments

The authors thank Prof. Jorg Grandl from Duke Neurobiology for providing the P1KO cell line and Piezo1 plasmid, and Dr. Kenneth Yamada from National Institutes of Health for the 16G3 and 13G12 antibodies. The authors thank Ashley Henderson for technical support, and Amanda Lewis for guidance in P1KO cell culture and Piezo1 plasmid transfection and helpful discussions. This work was supported by the National Institutes of Health Grant 5R37-DK052985-23.

## The table of contents

Microbubble-induced impulsive shear flow is involved in ultrasound therapies, resulting in Ca^2+^ signaling and cell injury. Here, treating cells with integrin-biding beads is demonstrated as an effective means to elicit Ca^2+^ signaling while avoiding cell damage. A novel signaling mechanism with increased integrin ligation to extracellular matrix, triggers ATP release and activation of P2X, but not Piezo1 ion channels, is discovered.

