## Supporting Information for "Mechanically Induced Integrin Ligation Mediates Intracellular Calcium Signaling with Single Pulsating Cavitation Bubbles"

### Supplementary Information Text

**Single bubble (SB)-generated shear flow: shear stress estimation at different γ.** Based on previous study with similar bubble dynamics, the shear stress τ near the substrate at fixed γ can be estimated from the velocity field [1]:

$$\tau\,(\mathrm{t}) = \rho\sqrt{\frac{v}{\pi}}\int_0^t \frac{\partial V(t\prime)}{\partial t\prime}\frac{dt\prime}{\sqrt{t-t\prime}}\ ,$$

which can be written in a discrete from:

$$\tau\,(\mathrm{t}_m) = \rho\sqrt{\frac{v}{\pi}}\left(\sum_{n=1}^{m}\frac{V_n-V_{n-1}}{\Delta t}\frac{\Delta t}{\sqrt{\mathrm{t}_m-t_{n-1}}}\right) = \rho\sqrt{\frac{v}{\pi}}\left(\sum_{n=1}^{m}\frac{V_n-V_{n-1}}{\sqrt{\mathrm{t}_m-t_{n-1}}}\right) \qquad [1]$$

where $\mathrm{t}_n$ and $V_n$ is the time and velocity for the recorded image frame # n, bubble is generated at frame n=0, $\Delta t$=0.5 μs, ρ is the density of the liquid medium (1000 kg $m^{-3}$), $v$ is the liquid kinematic viscosity (0.896 x $10^{-6}$ $m^2$ $s^{-1}$)

Moreover, considering the impulsive nature of the shear flow induced by single bubbles, we calculated shear stress integral (SSI) that incorporates the contribution of both the amplitude and the duration of the shear stress by:

$$SSI = \int_{t1}^{t2} \tau^{\beta} dt, \qquad [2]$$

Where t is the time, and $t_1$ and $t_2$ delineate the lower and upper integration limits during the shear flow. Previous studies have indicated that strain or stress integral may be appropriate for gauging the membrane poration under dynamic shear stress with a value of β ~ 2 [2-4]. The estimated shear stress is shown in Figure S2.

**General observation of cell spread area alteration with TB induced $Ca^{2+}$ response.** Transient increase, reduction and recovery of cell spread area are also observed in TB treated HeLa cells that show non-injury $Ca^{2+}$ response from our previous study [5], see Figure S4 (A-B). The results again suggest that $Ca^{2+}$ response is initiated during the transient increase of cell spread area following TB treatment (see the enlarged view). The intracellular $Ca^{2+}$ elevation is followed by cell spread area reduction (see the green dashed lines that reach the peak value successively), and the intracellular $Ca^{2+}$ decay preceded the cell spread area recovery. These results suggest $Ca^{2+}$ response is driving reduction and recovery of the cell spread area.

We found a similar trend of $Ca^{2+}$ response driving the reduction and recovery of the cell spread area in TB elicited injury $Ca^{2+}$ response w/o beads, see Figure S4C. Interestingly, for these injury $Ca^{2+}$ response, there is no obvious increase of the cell spreading area following the TB treatment, consistent with our observations for SB treated HEK293T cells with injury (Figure 5F).

A summary of the maximum cell spread area reduction in relation to the amplitude of the $Ca^{2+}$ response is shown in Figure S4D. With increased amplitude of $Ca^{2+}$ signaling, the individual cell's peak spread area reduction gradually transits from no $Ca^{2+}$ response (black symbols) to non-injury $Ca^{2+}$ response (blue symbols), and finally injury $Ca^{2+}$ response (red symbols). The data also reveals that the peak spread area reduction is below 5% for all the no $Ca^{2+}$ response cases, within 5-12% for the majority of the non-injury $Ca^{2+}$ response cases, and above 12% for most of the injury $Ca^{2+}$ response cases, similar to the results for SB treated HEK293T P1KO cells shown in Figure 5G.

**Error analysis for calcium response amplitude and cell spread area reduction.** The error in the normalized calcium response amplitude $\boldsymbol{f}$ and cell spread area reduction $\boldsymbol{A}$ is calculated by error propagation of

$$\boldsymbol{f} = \frac{\boldsymbol{F_p - F_0}}{\boldsymbol{F_0}} = \frac{\boldsymbol{F_p}}{\boldsymbol{F_0}} - \mathbf{1}\ , \quad \boldsymbol{A} = \frac{\boldsymbol{S_0 - S_p}}{\boldsymbol{S_0}} = \mathbf{1} - \frac{\boldsymbol{S_p}}{\boldsymbol{S_0}}$$

The uncertainty, where the related variables are baseline and peak value of fura2 ratio $\boldsymbol{F_0}$, $\boldsymbol{F_p}$, and spread area of the cell $\boldsymbol{S_0}$, $\boldsymbol{S_p}$. Let $\boldsymbol{u}$ denotes the uncertainty of the respective variables, the uncertainty of $\boldsymbol{f}$ can be derived as:

$$\boldsymbol{u_f} = \sqrt{\left(\frac{\partial \boldsymbol{f}}{\partial \boldsymbol{F_p}}\right)^2 \boldsymbol{u_{F_p}}^2 + \left(\frac{\partial \boldsymbol{f}}{\partial \boldsymbol{F_0}}\right)^2 \boldsymbol{u_{F_0}}^2} = \sqrt{\left(\frac{1}{\boldsymbol{F_0}}\right)^2 \ \boldsymbol{u_{F_p}}^2 + \left(\frac{-\boldsymbol{F_p}}{\boldsymbol{F_0}^2}\right)^2 \boldsymbol{u_{F_0}}^2} = \frac{1}{\boldsymbol{F_0}} \sqrt{\boldsymbol{u_{F_p}}^2 + \left(\frac{\boldsymbol{F_p}}{\boldsymbol{F_0}}\right)^2 \boldsymbol{u_{F_0}}^2}$$

As $\boldsymbol{u_{F_p}} \sim \boldsymbol{u_{F_0}}$ & $\frac{\boldsymbol{F_p}}{\boldsymbol{F_0}} = \boldsymbol{f} + \mathbf{1}$

$$\boldsymbol{u_f} = \frac{\boldsymbol{u_{F_0}}}{\boldsymbol{F_0}} \sqrt{\mathbf{1} + (\boldsymbol{f} + \mathbf{1})^2} \qquad [3]$$

Similarly, the uncertainty of $\boldsymbol{A}$:

$$\boldsymbol{u_A} = \frac{\boldsymbol{u_{S_0}}}{\boldsymbol{S_0}} \sqrt{\mathbf{1} + (\mathbf{1} - \boldsymbol{A})^2} \qquad [4]$$

$\boldsymbol{F_0}$ and $\boldsymbol{S_0}$ are the averaged value of the baseline, their SD and SEM can be easily calculated. Therefore, using eq.[3] and eq.[4] we can find the SEM of the normalized calcium response amplitude $\boldsymbol{f}$ and cell spread area reduction $\boldsymbol{A}$. The estimated error are shown in Figure 5, Figure S4D and Figure S5.

**Summary on the activation of piezo1 from previous studies using ultrasound or shear flow.** Compared to the reported activation of Piezo1 by ultrasound or shear flow in these listed studies where the stimulation duration is mainly from hundreds of millisecond to tens of second (Table S1), the duration of the produced shear flow from our single pulse cavitation bubble is much shorter (~ 20 µs).

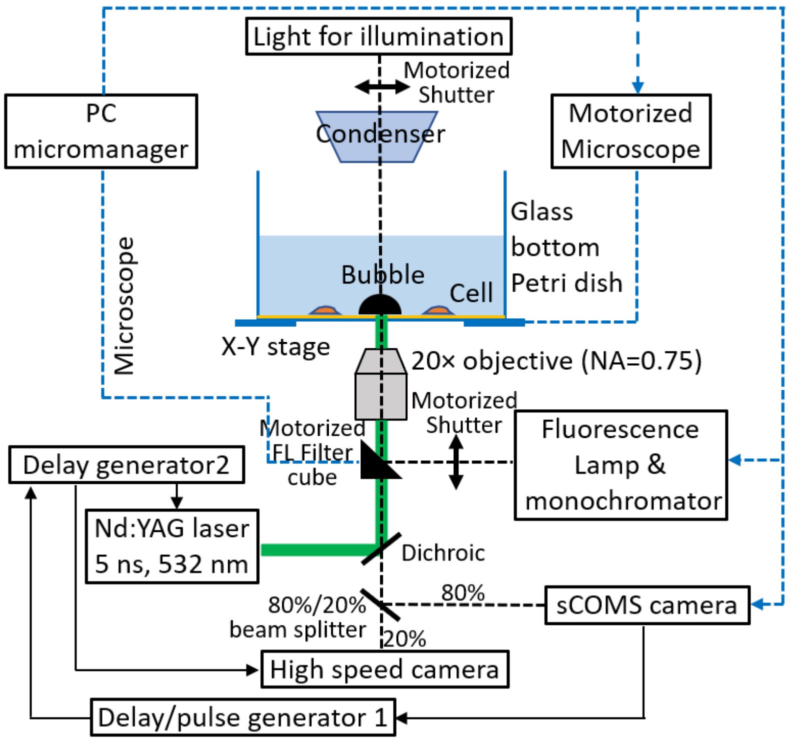


**Figure S1.** Experimental setup. Micromanager is utilized to synchronize the pulse laser (for SB generation), motorized microscope system (Axio Observer Z1; Zeiss), fluorescence monochromator system (DeltaRAM X; PTI) and cameras (high speed camera and sCOMS camera) for imaging and data acquisition. The exposure out signal of the sCOMS camera (EDGE 5.5 CL; PCO) is used to trigger the delay / pulse generator 1, which controls the delay generator 2 at 10 s later to give command for the high-speed image acquisition and pulse laser firing. The fluorescence (FL) excitation, corresponding FL filter cube, and image recording were automatically switched between fura-2 and PI with a switching interval about 0.7s through μManager, which communicates and coordinates the operation sequence between the sCOMS camera, the microscope, the fluorescence monochromator, and the motorized shutters for bright field and fluorescence imaging, respectively. For more details, see Materials & Methods (Image Recording and Analysis) in the main text.

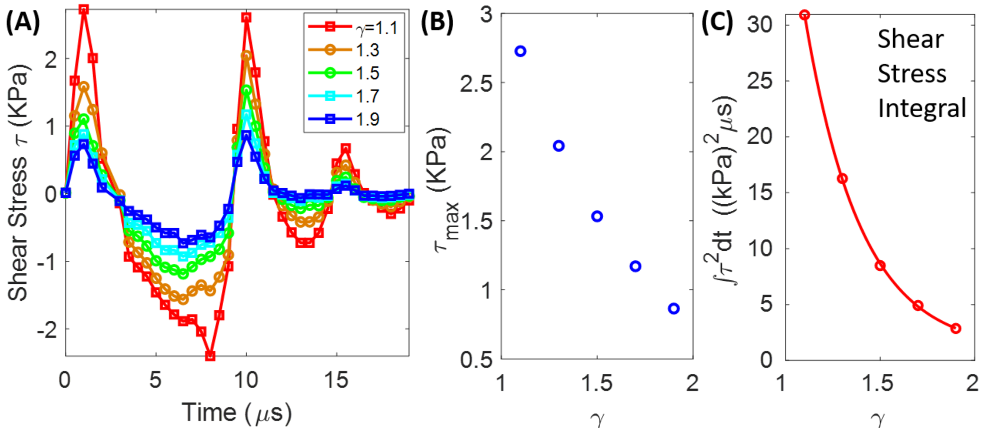


**Figure S2.** Estimation of SB-generated shear stress near the substrate using the PIV data. (A) Time evolution of the shear stress $\tau$ at different normalized standoff distance $\gamma$ using Eq.1. (B) The peak shear stress $\tau_{max}$ obtained from (A) vs $\gamma$. (C) Shear stress integral with time ($\int \tau^2 \mathrm{d}t$) vs $\gamma$. The sold line is an exponential fit to the data points: $y=1228\ e^{(-3.37x)} + 0.84$, with $R^2 = 0.99$.

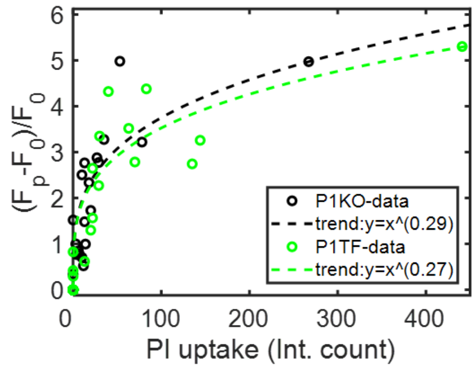


| PIKO ($\gamma$=1-1.5) | Injury $Ca^{2+}$ response | non-injury $Ca^{2+}$ response | No $Ca^{2+}$ response |
|---|---|---|---|
| N | 19 | 2 | 15 |
| Percentage (%) | 53 | 5 | 42 |
| PITF ($\gamma$=1-1.5) | Injury $Ca^{2+}$ response | non-injury $Ca^{2+}$ response | No $Ca^{2+}$ response |
| N | 13 | 2 | 17 |
| Percentage (%) | 41 | 6 | 53 |

**Figure S3.** Dependence of SB elicited intracellular $Ca^{2+}$ response on PI uptake and $\gamma$ in P1KO and P1TF cells. *Left*: Correlation of $Ca^{2+}$ response with PI uptake for the population of individual cells in the group of P1KO and P1TF among $\gamma$=1-1.5. Symbols represent data points measured from experiment while the dashed lines are power law trend of the data points, with $R^2$ of 0.9866 and 0.9949 for P1KO and P1TF, respectively. *Right*: a table showing the overall percentage for the three different kinds of cell response: $Ca^{2+}$ response with injury, $Ca^{2+}$ response without injury, no $Ca^{2+}$ response and no injury in the range of $\gamma$=1-1.5.

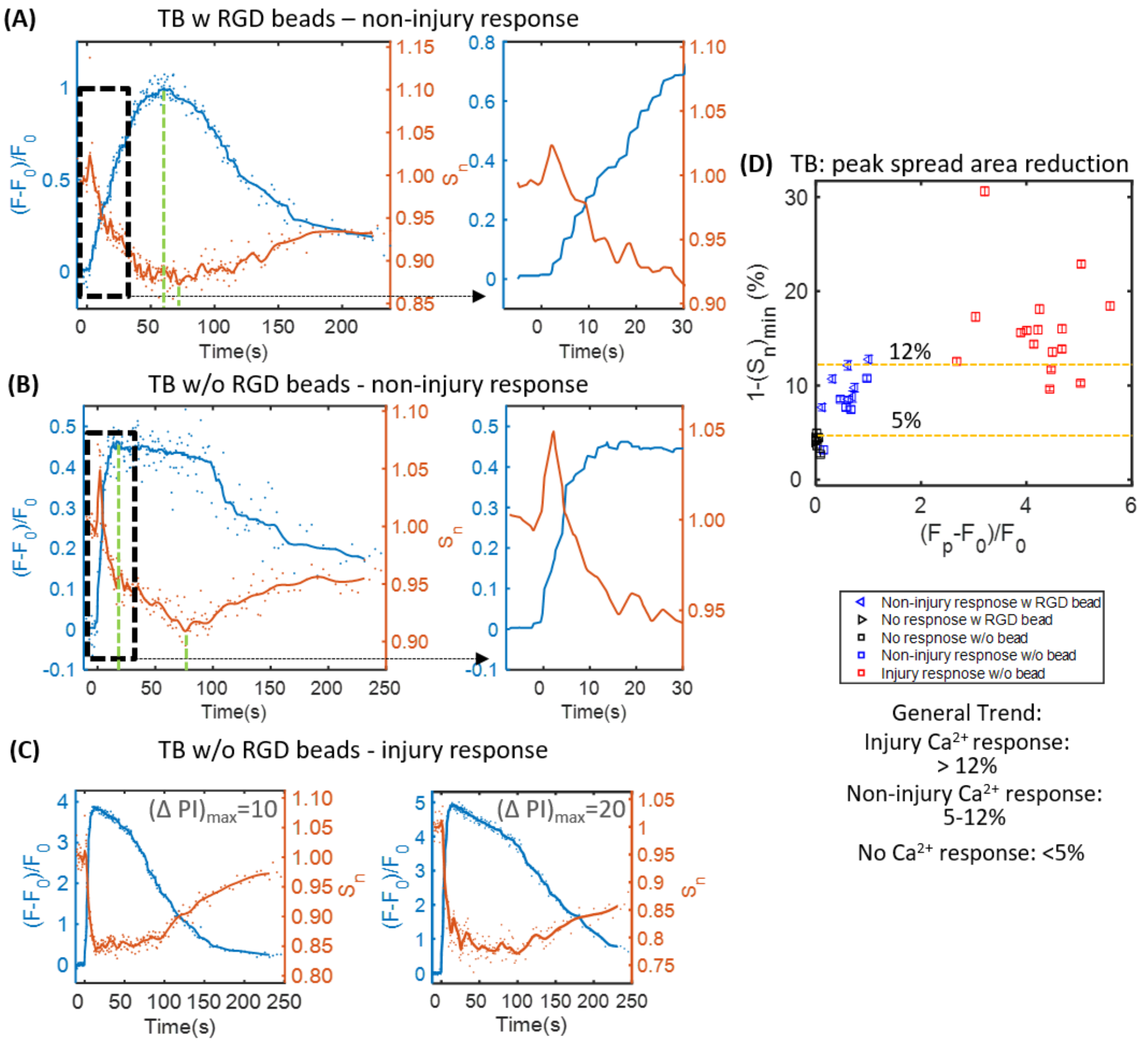


**Figure S4.** Transient increase and reduction of cell spread area for TB elicited different kinds of $Ca^{2+}$ response in HeLa cells from our previous work [5]. (A) A representative HeLa cell with RGD beads treated by TB at $S_d$=60 μm. (B) A representative HeLa cell without RGD beads treated by TB at $S_d$=50 μm. *Left:* time traces of intracellular $Ca^{2+}$ signaling (*blue*) and cell spread area alteration (*orange*). *Right*: enlarged view for the dashed box region on *left*, showing $Ca^{2+}$ response is initiated during cell spread area increase immediately following TB treatment. (C) Time traces of intracellular $Ca^{2+}$ signaling (*blue*) and cell spread area alteration (*orange*) for TB elicited injury $Ca^{2+}$ response in two representative HeLa cells w/o beads and with different amount of PI uptake ($(\Delta PI)_{max}$), respectively. (D) Summary of the maximum cell spread cell reduction for TB evoked different kinds of $Ca^{2+}$ response, either with RGD beads (*triangles*) or without beads (*squares*). The data shows that most cases of the injury $Ca^{2+}$ response, non-injury $Ca^{2+}$ response and no $Ca^{2+}$ response has a maximum spread area reduction of above 12%, between 5% and 12%, and below 5%, respectively. The error bars depict the SEM.

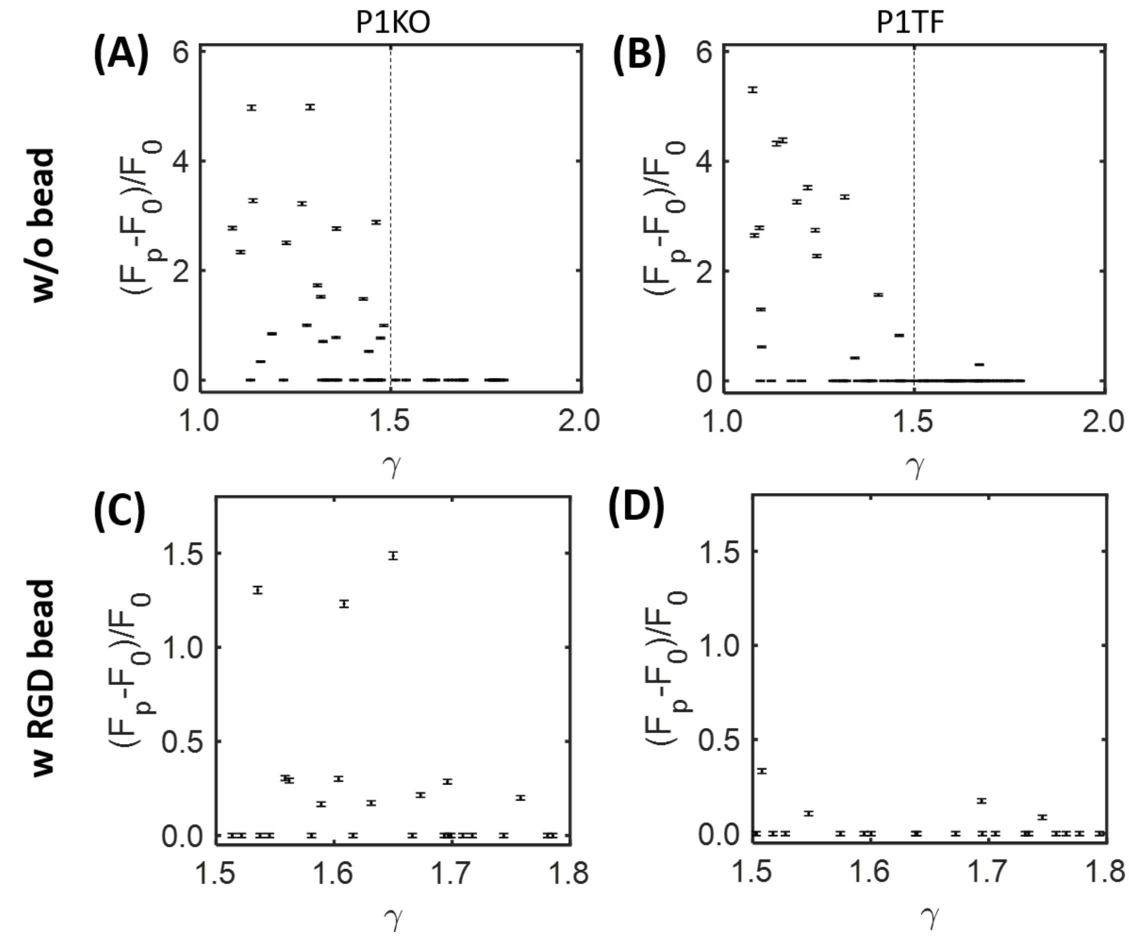


**Figure S5.** Error estimation for SB induced $Ca^{2+}$ response amplitude. (A) and (B) HEK293T P1KO and P1TF cells not treated with beads, corresponding to Figure 2F and Figure 2G in the main text, respectively. (C) and (D) HEK293T P1KO and P1TF cells treated with RGD coated beads, corresponding to the left and right panels in Figure 3E, respectively. The error bars show the SEM of the data.

**Table S1.** Summary on the activation of piezo1 from previous studies using ultrasound or shear flow.

| Previous studies | Stimuli type | **f (MHz)** | Intensity or pressure | **Total treatment time** | PRF | duty cycle | Single pulse duration | Flow velocity and shear stress | cell system | Detection method for piezo1 activation | results |
|---|---|---|---|---|---|---|---|---|---|---|---|
| Prieto, M. L., et al. (2018) [6] | Continuous US | 43 | 50 or 90 $W/cm^2$ | **200 ms** | N.A. | N.A. | N.A. | Acoustic streaming velocity <0.14 mm/s | HEK293T transfected with Piezo1 | Electrophysiological Recording (patch-clamp) | Piezo1 is activated |
| Pan, Y., et al. (2018) [7] | US | 2 | ~0.6 MPa | **5 s** | 5Hz | 10% | 20ms | N.A. | HEK293T transfected w Piezo1 | $Ca^{2+}$ imaging | cannot activate Piezo1 unless with integrin-binding microbubble |
| Qiu, Z., et al. (2019)[8] | US | 0.5 | 0.3 MPa | **200 ms** | 1000 Hz | 40% | 400 µs | N.A | HEK293T transfected w Piezo1 | $Ca^{2+}$ imaging | Piezo1 can be activated |
| D.F. Liao, et al. (2019)[9] | VD-SAW | 30 | ~1.6MPa | **60 s** | 2Hz | 20% | 100ms | Streaming velocity of 0.8m/s, shear stress 50 $dyne/cm^2$ | HEK293T Piezo1 KO and transfected | $Ca^{2+}$ imaging | Piezo1 can be activated |
| | | 30 | ~1.6MPa | **60 s** | 200Hz | 20% | 1ms | Streaming velocity of 0.8m/s, shear stress 50 $dyne/cm^2$ | HEK293T Piezo1 KO and transfected | $Ca^{2+}$ imaging | Piezo1 has no significant effect |
| Ranade, S. S., et al. (2014)[10] | Perfusion shear flow | Single pulse | N.A. | **600ms** | N.A. | N.A. | 600 ms | Average velocity of 6.9-21 mm/s, shear stress 23.3-70.2 $dyne/cm^2$ | HEK293T transfected w Piezo1 | Electrophysiological Recording (patch-clamp) | Piezo1 can be activated |
| This study | Laser-induced bubble | Single pulse | Bubble radius 45-55 µm | **~20 µs** | Single pulse | N.A. | ~20 µs | Maximum velocity of 1.6-2.5 m/s ($\gamma$: 1.5-1.9); maximum shear stress 0.9-1.5 kPa (9000-15000 $dyne/cm^2$) | HEK293T Piezo1 KO and transfected | $Ca^{2+}$ imaging | Piezo1 has no significant effect |
